## Supplemental figures for "Genome-wide antibiotic-CRISPRi profiling identifies LiaR activation as a strategy to resensitize fluoroquinolone-resistant *Streptococcus pneumoniae*"

### Supplementary figures

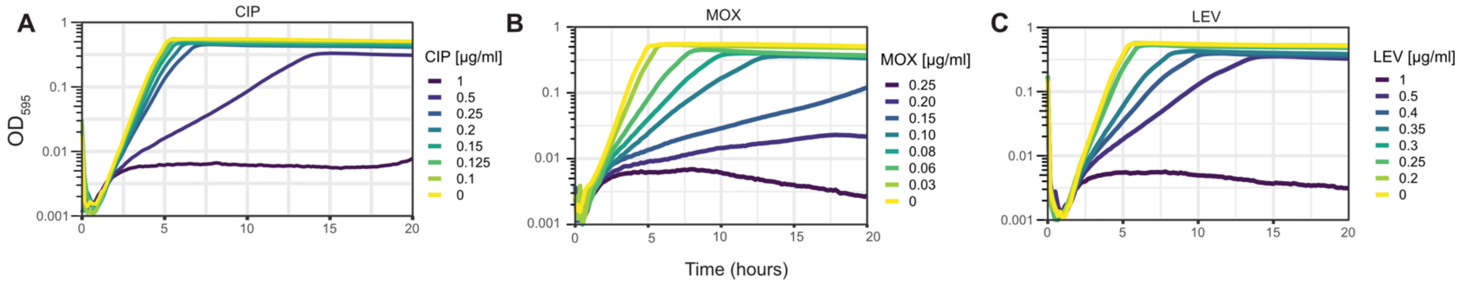


**Supp Fig 1. Growth curves of the *S. pneumoniae* CRISPRi library treated with a range of antibiotic concentrations.** The *S. pneumoniae* CRISPRi library was treated with a panel of fluoroquinolones at varying concentrations to determine the sub-lethal dose. Growth curves are shown here for **A)** ciprofloxacin (CIP) **B)** moxifloxacin (MOX) and **C)** levofloxacin (LEV). Sub-lethal concentrations used for the screens were 0.5, 0.06, and 0.35 µg/ml respectively.

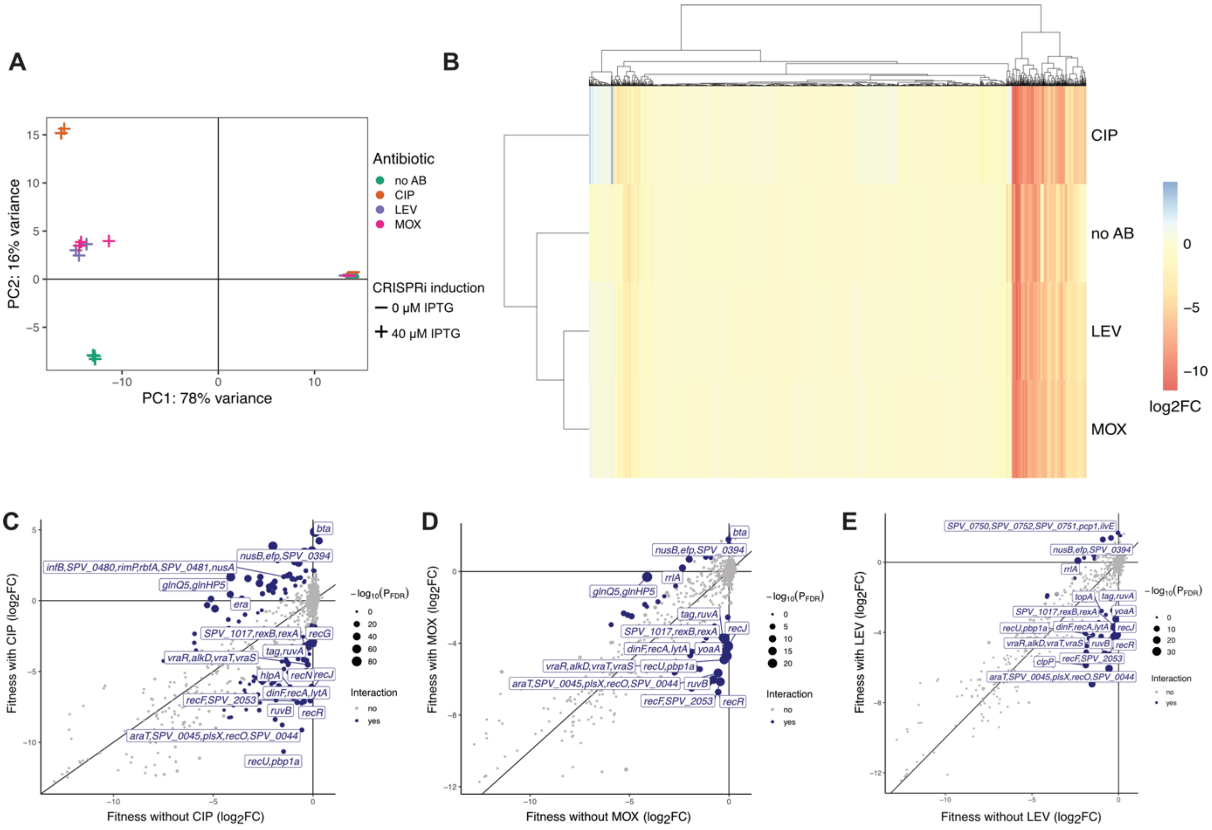


**Supp Fig 2. CRISPRi-seq screen with *S. pneumoniae* library treated with fluoroquinolones. (A)** Principal component analysis served as a quality control measure and indicated distinct clusters between induced and uninduced samples, as well as between antibiotic treated and untreated samples (no AB). **(B)** Heatmap depicting scaled fitness scores of sgRNA targets with a significant differential fitness. This provides a broad view of gene essentiality with no antibiotic treatment and with antibiotic treatment. A negative score (red) indicates a fitness loss upon gene knockdown and a positive score (blue) indicates a fitness gain compared to no antibiotic. Conditional essentiality scatter plots for each fluoroquinolone screen **C)** ciprofloxacin, **D)** moxifloxacin, **E)** levofloxacin. Screens were conducted *in vitro* in C+Y medium. Fitness scores are measured by the log2 fold change of sgRNA counts between induced and uninduced samples. Points in blue indicate a significant interaction between induction and antibiotic treatment and points in grey indicates a non-significant interaction. Point size indicates the -log10 transformed *p*-adj value.

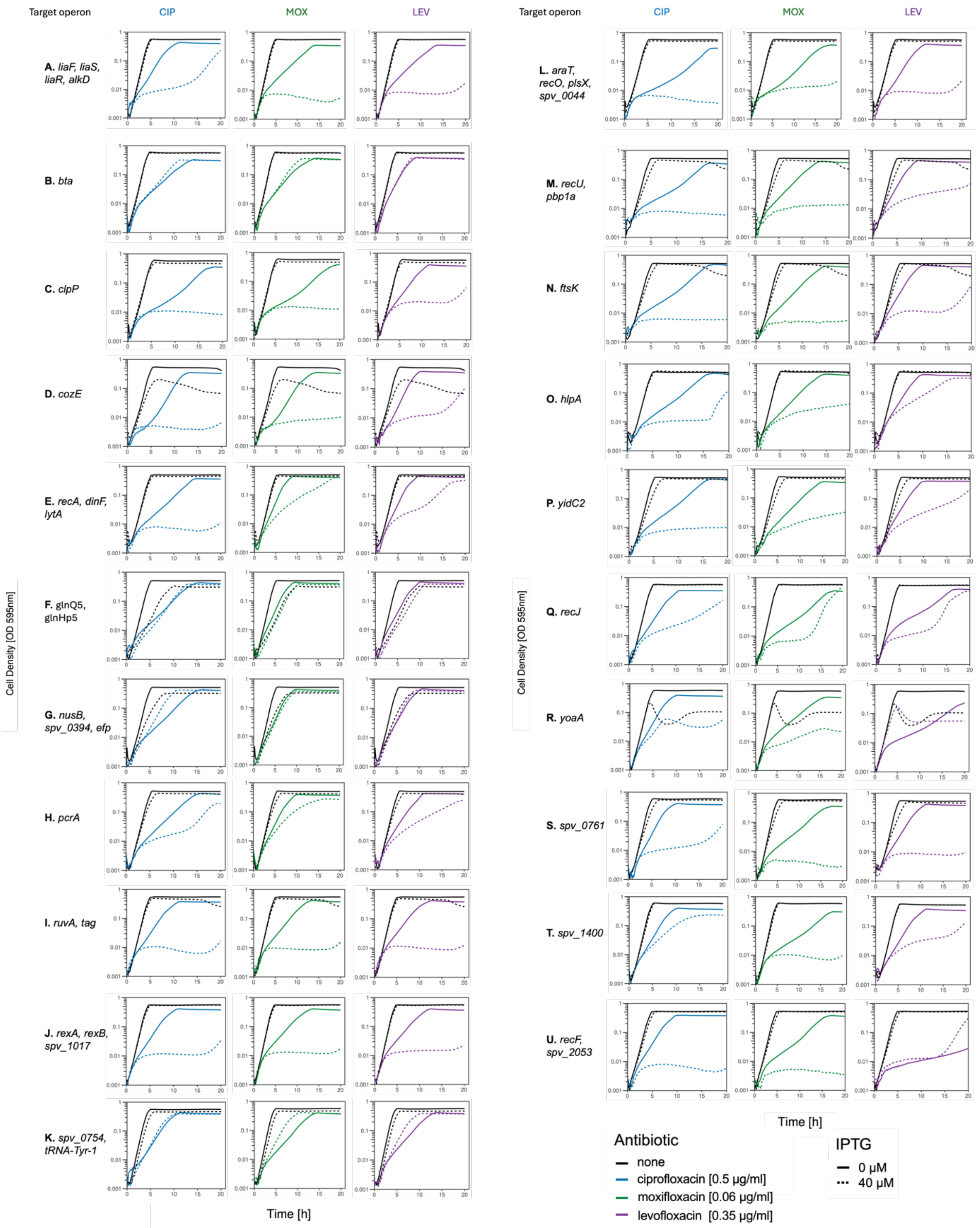


**Supp Fig 3. Growth profiles of CRISPRi strains treated with fluoroquinolones. (A-U)** Individual CRISPRi strains were made for 21 sgRNAs that were some of the top hits from the CRISPRi-seq screens. Strains were treated with CIP, MOX and LEV at sub-lethal concentrations to determine if the growth phenotype of the single mutants could be validated based on the results of the pooled CRISPRi-seq screen.

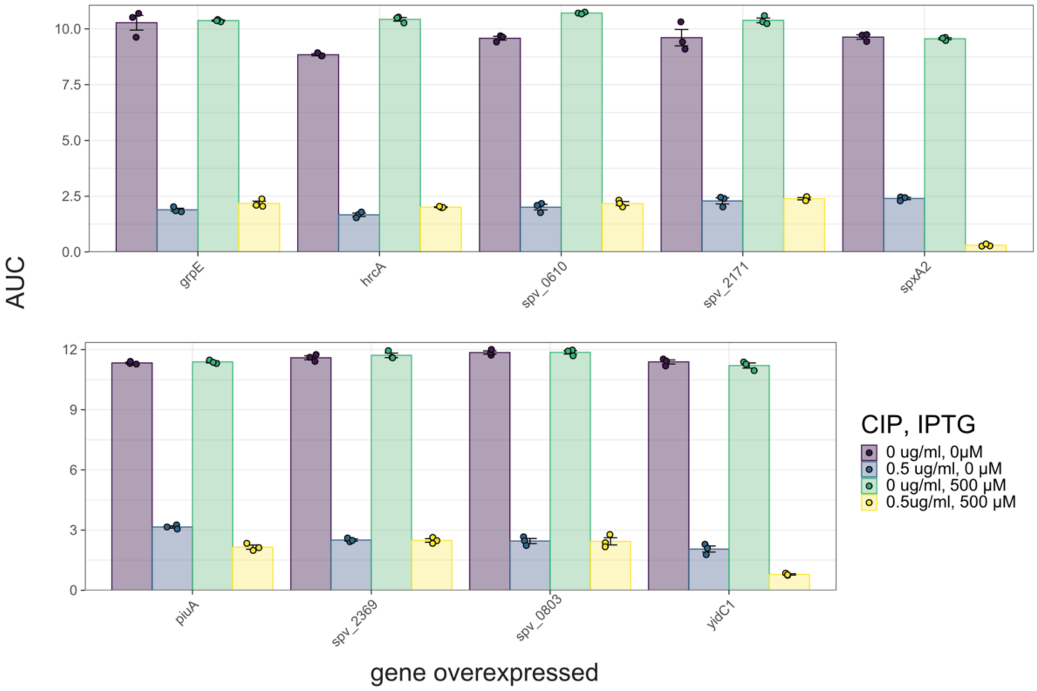


**Supp Fig 4. Area under the curve (AUC) plots for overexpression strains.** Overexpression strains were generated by placing a second copy of the gene at an ectopic locus under the control of $p_{lac}$ and inducing with IPTG. Strains were treated with a sub-lethal concentration of CIP and effect on growth was assessed for 20 hours.

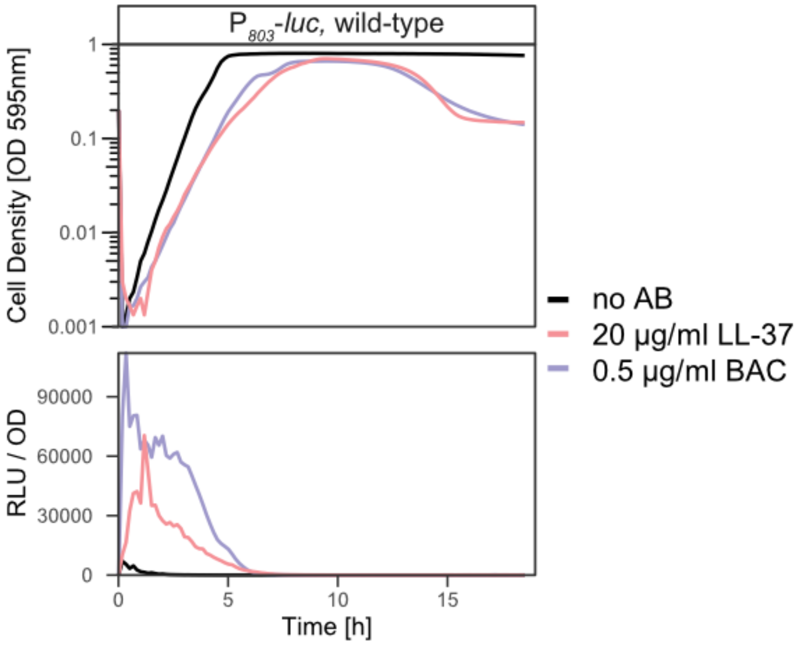


**Supp fig 5. LL-37 induced the LiaR regulon.** In addition to bacitracin, the antimicrobial peptide LL-37 also induced expression of luciferase from the strain $P_{spv\_0803}$-*luc*, which served as a proxy for LiaR induction.

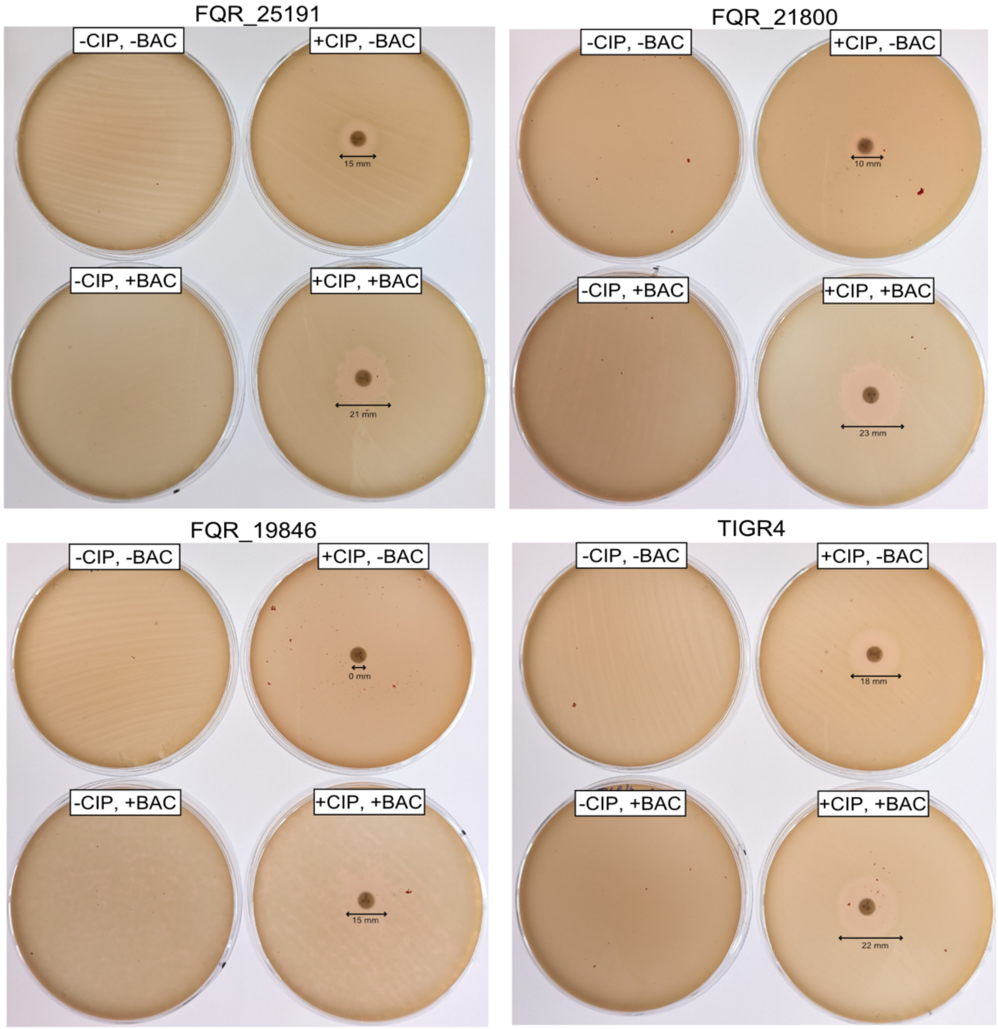


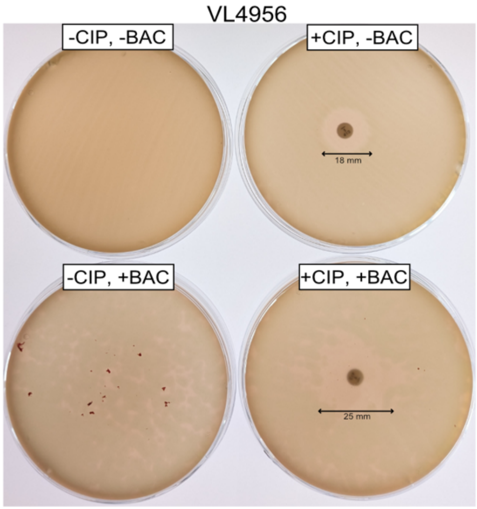


| Strain | Isolate description |
|---|---|
| FQR_25191 | $LEV^R$ clinical isolate from sputum, serotype 31 |
| FQR_21800 | $LEV^R$ clinical isolate from sputum, serotype 19F |
| FQR_19846 | $LEV^R$ clinical isolate from trachea, serotype 9V |
| TIGR4 | Lab strain, serotype 4 |
| VL4956 | D39V, $CIP^R$ *parC* S79F |

**Supp Fig 6. Ciprofloxacin and bacitracin synergize against fluoroquinolone resistant clinical pneumococcal strains.** Ciprofloxacin disc diffusion assays show larger zones of inhibition in combination with bacitracin for several strains of different serotypes.

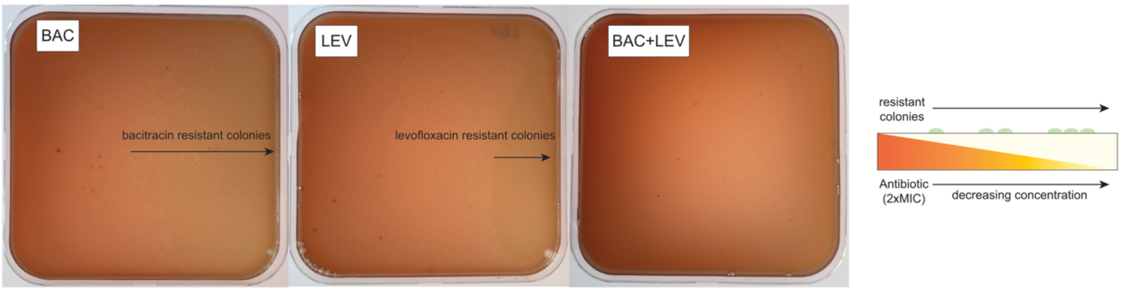


**Supp Fig 7. No resistant colonies were observed when WT *S. pneumoniae* was exposed to a combination of bacitracin and levofloxacin.** Concentration gradient plates show that resistant colonies could be obtained for bacitracin and levofloxacin individually at 2×MIC concentrations. However, no resistant colonies were obtained when bacitracin and levofloxacin were in combination.

A

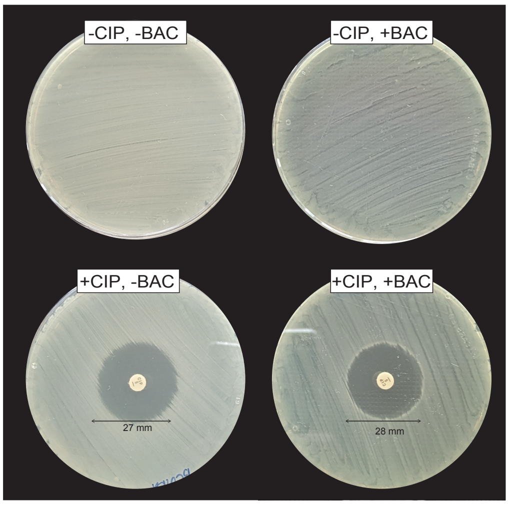


B

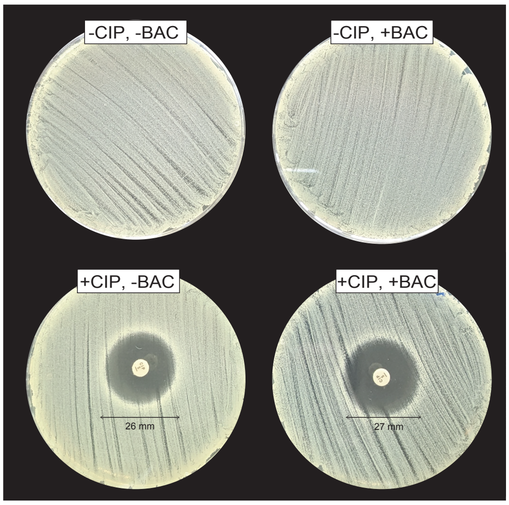


**Supp Fig 8. Ciprofloxacin and bacitracin do not synergize against two other Gram-positive species.** Ciprofloxacin disc diffusion assays for **A)** *B. subtilis* and **B)** *S. aureus*. There was no difference in the zone of inhibition between ciprofloxacin alone and when combined with bacitracin.

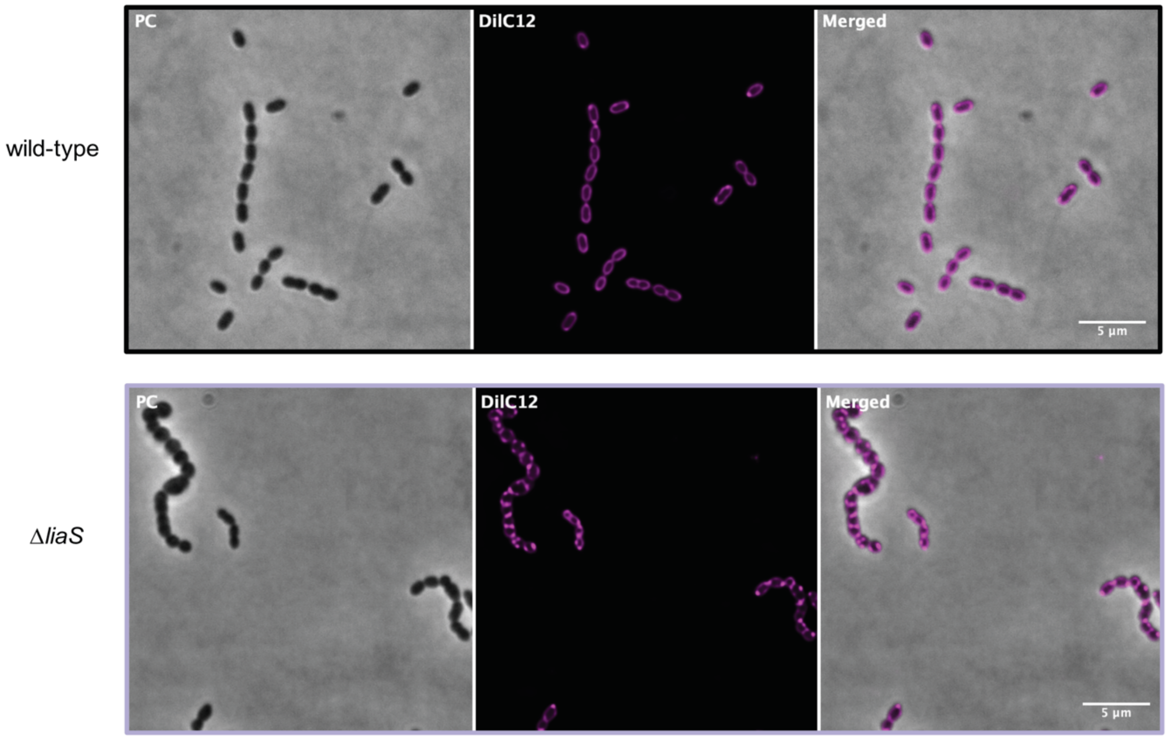


**Supp Fig 9. The Δ*liaS* mutant displays perturbed cell membrane physiology compared to WT, as visualized by DilC12 staining.** DiI-C12 dye incorporates into fluid areas of the cell membrane lipid bilayer, thus allowing visualization of cell membrane integrity. The Δ*liaS* mutant cells displayed irregular cell morphologies with either swollen or shrunken cells. Patches of fluorescent foci could be observed due to the accumulation of DiI-C12 to these highly fluid lipid regions, indicating disruption of the cell membrane
